# A Non-invasive, Biomarker Assay for Detecting Chronic Wasting Disease Pathology in White-tailed Deer

**DOI:** 10.1101/2022.07.20.500695

**Authors:** Robert D. Bradley, Emma K. Roberts, Asha E. Worsham, Megan N. Ashton, Emily A. Wright, Ned Saleh, Daniel M. Hardy

**Affiliations:** Department of Biological Sciences, Texas Tech University, 2901 Main Street, Lubbock, TX 79409; Natural Science Research Laboratory at the Museum of Texas Tech University, 3301 4^th^ Street, Lubbock, TX 79415; Climate Sciences Center, Texas Tech University, 2500 Broadway Ave, Lubbock, TX 79409; Department of Cell Biology & Biochemistry, Texas Tech University Health Sciences Center, 3601 4^th^ St., Lubbock, TX 79430; Convergent Animal Health LLC, 118 Thoroughman Ave., Ferguson, MO 63135

## Abstract

We describe a blood test that exploits differences in the abundance of diagnostic miRNA biomarkers associated with prion infections in cervids. Using sera from 93 pen-raised white-tailed deer euthanized after unintentional exposure to Chronic Wasting Disease (CWD), quantification of candidate reference and diagnostic miRNAs revealed sensitivity of the q-RT-PCR method to interference from sample degradation, requiring exclusion of 60 specimens that exhibited excessive hemolysis or yielded poor amplification of reference or diagnostic miRNAs. Subsequent quantification of three potentially diagnostic and two control miRNAs in the 33 remaining, minimally degraded sera established diagnostic criteria congruent (100% sensitivity, 92.3% specificity, 93.9% accuracy) with results from standard CWD diagnosis by microscopic detection of immunoreactivity in obex and/or medial retropharyngeal lymph nodes (conducted by the National Veterinary Services Laboratory, NVSL, Ames, IA). Overall, the miRNA assay proved to be at least as accurate and sensitive as other CWD testing alternatives to immunohistochemical diagnosis, albeit supported by data from relatively few animals owing to the challenge of acquiring usable specimens from animals euthanized in a mass depopulation. Nevertheless, for sera acquired antemortem, this biomarker-based test may represent a useful new resource for managing spread of CWD that offers the advantages of being rapid, sensitive, non-invasive, and amenable to high throughput scaling.

## INTRODUCTION

Chronic Wasting Disease (CWD) is a transmissible spongiform encephalopathy (TSE), associated with the deer family Cervidae (Williams and Young 1980; Hannaoui et al. 2017; Slota et al. 2019), that is caused by alternative folding of a prion protein into a pathogenic conformation (*PrP*^*CWD*^, Prusiner 1998; Whitechurch et al. 2017; Marín-Moreno et al. 2017; Li et al. 2021). Pathologically folded PrP^CWD^ typically forms amyloid plaques that aggregate predominantly in the central nervous system and lymphoid tissues. Over time, this infection results in a continual decline in the animal’s health, invariably leading to death. CWD is related to other prion diseases (Saá et al. 2006) such as those associated with cattle and bison (*Bos taurus* and *B. bison*; bovine spongiform encephalopathy), sheep (*Ovis aries*; scrapie), and humans (*Homo sapiens*; Creutzfeldt-Jakob disease and kuru). To date, CWD is the only prion disease to be documented in both captive and wild mammals (Sohn et al. 2002; Benestad et al. 2016; Kramm et al. 2017; Vikøren 2019; USGS 2021) and has been shown to be highly contagious within and among most cervid species. As with most TSEs, it is thought that CWD is not easily, if at all, transmissible to humans (Osterholm et al. 2019; Barria et al. 2018), although recent studies provide evidence that CWD transmission could occur across the human (or livestock) barrier (see Waddell et al. 2017, Otero et al. 2021, Pritzkow et al. 2022).

CWD was first identified in captive mule deer (*Odocoileus hemionus*) from Colorado in 1967 (Williams and Young 1980). The disease range has since expanded to include 30 U.S. states, four Canadian provinces, and four foreign countries (Finland, Sweden, Norway, and South Korea; Osterholm et al. 2019; Slota et al. 2019). CWD has also been detected in captive cervid populations in 18 U.S. states and three Canadian provinces (https://www.usgs.gov/centers/nwhc/science/expanding-distribution-chronic-wasting-disease). Current evidence (Falcão et al. 2017; Mawdsley 2020; Buchholz et al. 2021; Hannaoui et al. 2017; Sohn et al. 2002; Hamir et al. 2011; Nalls et al 2013; Benestad et al. 2016; Vikøren 2019; Cullingham et al. 2020; USGS 2021) indicates that CWD occurs in several native North American cervid species, including white-tailed deer (*O. virginianus*), North American elk (*Cervus canadensis*), and moose (*Alces alces*); non-North American species such as reindeer (*Rangifer tarandus*), sika deer (*C. nippon*), red deer (*C. elaphus*), Reeve’s muntjac (*Muntiacus reevesi*), and fallow deer (*Dama dama*); and non-cervid species such as mink (*Vison vison*), ferrets (*Mustela furo*), feral and domestic cats (*Felis catus*), feral and domestic sheep, goats (*Capra hircus*), cows, pigs (*Sus scrofa*), and squirrel monkeys (*Saimiri spp*.). The species promiscuity of CWD is remarkable considering the allelic variation just among individual elk (O’Rourke et al. 1999), reindeer (Güere et al. 2019), or white-tailed deer (Seabury et al. 2020) influences disease susceptibility

CWD transmission can occur by numerous routes. Several studies indicate that CWD may be transmitted through consumption of carcasses and antlers (bone and velvet), tissues, saliva, feces, urine, blood, placenta, and semen (Mathiason et al. 2006; Haley et al. 2009, 2011; Tamgüney et al. 2009; Maddison et al. 2010; Terry et al. 2011; Kramm et al. 2019; Mysterud et al. 2020). Infected, asymptomatic individuals can shed large volumes of prions (Mathiason et al. 2009); further, prion shedding can occur several months prior to the visually characteristic “wasting phase” of the disease. Prions also have been shown to remain in soils for several years (Johnson et al. 2006, 2007; Seidel et al. 2007; Dorak et al. 2017; Gemovesi et al. 2007) allowing CWD and other TSEs to persist under multiple environmental conditions. CWD prions can also be taken up by grasses (Pritzkow et al. 2015) and sequestered in stems and leaves, thereby serving as a potential vector for horizontal transfer among herbivore species. Likewise, mineral licks (salt blocks for cattle) in Wisconsin are thought to serve as reservoirs for CWD and potentially represent a conduit for intra- and inter-species infection (Plummer et al. 2018). Even airborne transmission was shown to be possible under controlled laboratory conditions (Denkers et al. 2013), signaling another potential mode for animal-to-animal transmission.

Although little evidence suggests that CWD can be naturally transmitted from cervids to non-cervid species (Belay et al. 2004; Vaske and Lyon 2011; Williams et at. 2018), given the sympatric distribution of cervids and livestock, the introduction of CWD or other potential variant prion disease would be catastrophic to the livestock industry. Free-ranging and captive populations of deer as well as other cervid species represent a multibillion-dollar industry (Li et al. 2021) that is important from an ecological, socio-cultural, and economical perspective. It is estimated that 5.8 million deer are harvested in the U.S. alone (National Shooting Sports Foundation 2017), not including nearly 100,000 white-tailed deer from captive populations in Texas. Consequently, CWD outbreaks would be financially devastating to landowners participating in the hunting and/or deer breeding industries as some states may require all individual animals in a breeding facility, pen or enclosure, or specific site be destroyed based on a positive test from a single animal. Proactive detection methods that are highly sensitive for CWD would allow hunters, breeders, landowners, and state and federal agencies to enact critical management policies that minimize the impact of this disease at both local and regional scales.

To date, several diagnostic tests have been developed for detecting prions in blood or tissue samples. Protein misfolding cyclic amplification (PMCA) was among the first techniques capable of detecting prions in blood (Saborio et al. 2001; Soto et al. 2002; Castilla et al. 2005). Immunochemical techniques, including immunohistochemistry (IHC) and enzyme linked immunosorbent assay (ELISA) (Hibler et al. 2003; Keane et al. 2008, 2009) led to increased detection rates, simplicity of testing, and the ability to test additional tissue types such as: brainstem (obex), medial retropharyngeal lymph node, muscle, tonsil, and rectal biopsy (Hibler et al. 2003; Daus et al., 2011; Saa et al 2006; Thomsen et al. 2012). Atarashi et al. (2007) reported the development of a new method, Real-Time Quaking-Induced Conversion (RT-QuIC), as a highly sensitive assay that amplifies CWD prions. Haley et al. (2014, 2016) and Holz et al. (2022) further developed this method and extended testing capabilities to include antemortem samples such as nasal swabs (effective only for end stage CWD) and rectal biopsies. This method successfully detected prions in skeletal muscles of infected white-tailed deer, leading Li et al. (2019) to promote RT-QuIC as a high-throughput method for monitoring hunter-harvested venison. Further, Christenson et al. (2022) produced a modified version of the RT-QuIC method (MN-QuIC™) that utilizes gold nanoparticles (AuNPs) to assist in increasing diagnostic sensitivity and to serve as a relatively quick and portable assay. Collectively, the available tests that detect prion deposition (IHC, ELISA) or activity (PMCA, RT-QuIC, MN-QuIC) provide useful tools for CWD diagnosis and surveillance, but none have yet proved to be effective for high throughput monitoring of infection and transmission in living animals.

Slota et al. (2019) used differential expression of miRNAs, small non-coding RNA transcripts approximately 18–22 nucleotides in length, as biomarkers for predicting the presence of CWD prions in elk (*C. canadensis*). These highly specific and sensitive biomarkers circulate in peripheral blood and show tremendous potential in predicting a wide range of diseases such as schizophrenia, diabetes, cancer, infectious and autoimmune diseases (Richens et al. 2016). Slota et al. (2019) reported that the abundance of 47 microRNAs (miRNAs) differed between 35 CWD positive elk and an equal number of non-infected individuals. Of those 47 variable miRNAs identified, 21 showed potential in producing diagnostic signatures correlated with prion infections in individual animals. Further, Slota et al. (2019) suggested that miRNA surveillance methods held promise at diagnosing sub-clinical prion infections that IHC and PMCA methods might miss.

Because CWD is a reportable disease, approved diagnostic assays are restricted to IHC and ELISA methods performed only by USDA-APHIS-approved and licensed State or Federal veterinary diagnostic laboratories. The biological specimens most often used for these tests must be obtained post-mortem (e.g., obex or retropharangeal lymph node). In the rare instances where antemortem testing is done, specimens are finite with respect to repetitive sampling (e.g., rectal lymph nodes), and are acquired by complex, invasive methods that require anesthesia and/or surgery performed by veterinarians or other highly skilled personnel. Furthermore, criteria for IHC diagnoses vary depending on the tissue examined (obex, retropharangeal lymph nodes, or rectal lymph nodes), and must be assessed by qualified pathologists.

The tremendous need for aggressive CWD surveillance worldwide requires new testing methods amenable to high throughput application. The main limitations of current tests include impossibility (e.g. obex=brain stem) or difficulty (e.g. rectal lymph nodes) in acquiring disease-affected tissues reliably and consistently by antemortem sampling. Here we applied rapidly emerging miRNA technologies to develop a less invasive, non-surgical, antemortem blood test that can be used for early detection of CWD infection in ongoing disease surveillance and management. The goal of this study was to ascertain the potential of using miRNA abundance in blood exosomes as an indicator of a prion pathology in individual cervids that presumably would lead to symptomatic CWD. Specifically, we identified differences in the abundance of miRNA biomarkers that correlate with CWD diagnosed by the standard IHC method approved for and used by state and federal agencies in their routine CWD surveillance, then devised a three-marker diagnostic protocol that reliably reflects CWD infection status. Finally, we comment on the power of using a consistent set of blood miRNAs for predicting presence of prions and ultimate risk of CWD.

## MATERIALS AND METHODS

### Sample Acquisition

In November 2021 the Texas Parks and Wildlife Department (TPWD) collected specimens from 300 pen-raised white-tailed deer euthanized, per TPWD and Texas Animal Health Commission regulations, because CWD had been detected in the herd, located at a breeding facility in Uvalde County, Texas. Animals were euthanized by the landowner and specimens obtained post-mortem as part of a TPWD-sponsored CWD research initiative. TPWD staff collected obex and medial retropharyngeal lymph nodes from all 300 individuals and sent them to NVSL (Ames, IA) for definitive CWD diagnosis by IHC, and collected blood and muscle samples from 168 of the 300 individuals for use in research to be conducted at our institution. Blood collected from the jugular vein or from pools within the neck near the decapitation area were placed into lavender K_2_ ethylenediamine-tetraacetic acid (EDTA) and red/grey serum separator (SST) Vacutainer® collection vials (Becton Dickinson Co., Franklin Lakes, NJ). Skeletal muscle specimens were either frozen in liquid nitrogen (later archived at -80° C), placed on ice (stored at either 4° C or -20° C and later archived at -80° C), or preserved in RNA*later* (Ambion, Inc., Austin, Texas) and archived at -80° C for future studies. All samples were deposited into the Natural Science Research Laboratory (NSRL; Texas Tech University, 2500 Broadway Lubbock, Texas 79409) where they were assigned a TK number (unique identifying number), placed in a barcoded cryotube, and stored at -80° C, in perpetuity, for future research studies. Corresponding data included locality, date of collection, sex, tissue type, sampling conditions, etc., and were incorporated into NSRL’s database.

### Sample Processing

Blood collection tubes were centrifuged 15 min at 1400 *g*, and sera drawn off for transport on ice to Texas Tech University where they were stored at -20° C, and later moved to -80° C. Among the 168 samples, 75 were immediately triaged by visual inspection for extreme hemolysis owing to the difficult field conditions under which they were acquired, leaving 93 specimens for initial miRNA studies.

### RNA Extraction and cDNA Synthesis

Total RNA was extracted from 200 uL of serum using the miRNeasy Serum/Plasma Kit (Qiagen Inc., Valencia, California, USA) per manufacturer’s instructions. First-strand cDNA synthesis was achieved using the miRCURY LNA RT kit (Qiagen Inc., Valencia, California, USA) also per manufacturer’s instructions, using an Eppendorf 5331 MasterCycler thermal cycler.

### Diagnostic miRNA Selection

Five miRNA primer sets (18–22 nucleotides in length; A-E) were evaluated following the protocols established by Convergent Animal Health (St. Louis, MO; patent application 22/63388262) to: 1) establish an endogenous control, independent of hemolysis and any known CWD association, for normalizing values of miRNA amplification (miRNA A), 2) independently assess extent of hemolysis (miRNA B), and 3) determine diagnostic potential of miRNAs as biomarkers for CWD prion pathology (miRNAs C, D, E).

### miRNA Amplification and Quantification

cDNA was amplified with the miRCURY LNA miRNA SYBR Green Polymerase Chain Reaction (PCR) Kit for Exosomes, Serum/Plasma, and other Biofluid Samples (Qiagen Inc., Valencia, California, USA). Assembled reactions in 96-well plates with optical adhesive covers were amplified using a 12K Flex Real-Time PCR thermal cycler (Life Technologies Inc., Carlsbad, CA) per manufacturer’s instructions, and crossing threshold values (Ct) of the reference and hemolysis control miRNAs and the five diagnostic miRNAs were recorded using the 12K Flex Real-Time PCR software. The relative abundance (ΔCt) of miRNAs B, C, D, and E were obtained by subtracting the mean Ct value (of experimental duplicates) for reference miRNA A from the mean Ct values obtained for the diagnostic and hemolysis control miRNAs (B, C, D, E).

### Hemolysis Assays

Given the large number of individual deer sampled, the post-mortem timeframe for sample collection ranged from 1-12 hours, with a mean of 3-4 hours, which could affect specimen integrity. To prioritize samples, we scored serum quality using three different hemolysis detection methods: 1) read count ratio >5 between miRNA B that is abundant in red blood cells (Slota et al. 2019) and miRNA A known to be unaffected by hemolysis (Shah et al. 2016; Slota et al. 2019); 2) spectrophotometrically (NanoDrop, ThermoFisher Scientific, Waltham MA), as hemoglobin-specific absorbance at 414 nm; and 3) visually, (with two independent raters) using the CDC color palette (https://www.cdc.gov/ncezid/dvbd/stories/research-lab-diagnostics/hemolysis-palette.html).

### Exclusion Criteria

Following the suggestions of Richens et al. (2016) for evaluating a diverse set of questions (i.e., effects of variation in sample acquisition on miRNA quantification, identification of the optimally diagnostic markers, etc.), three increasingly strict exclusion criteria were established. The first criterion, applied to all samples, was no prior thawing, as freeze-thaw is known to perturb miRNA quantification (Glinge et al 2017). The second and third criteria were applied sequentially to assess anticipated variation in quality of samples owing to differences in their acquisition. The second criterion excluded samples with a hemoglobin concentration <1.2 mg/ml (Kosecki et al. 2021), assessed visually and/or spectrophotometrically as described above. And the third criterion excluded samples with amplification of reference miRNA A with Ct > 28, which served as a reference for overall miRNA quantities in the samples. Applying all three exclusion criteria left N=33 minimally degraded sera (hemoglobin concentration <1.2 mg/ml and robust amplification of miRNA A), which we designated as the “Group I” dataset. Then, for downstream analyses assessing validity of the exclusion criteria, we captured additional samples by progressively relaxing the criteria. “Group II” comprised the 33 sera from Group I plus six sera that yielded poor amplification of miRNA (i.e., third exclusion criterion not applied; N=39). And “Group III” comprised the 39 sera from Group II plus 32 sera with hemoglobin concentration >1.2 mg/ml (i.e. neither second nor third exclusion criteria applied) that had also returned Ct values for all five miRNAs (N=71).

### Statistical Analysis

Discriminant Function Analysis was performed on Groups I-III using JMP Pro software (SAS Institute Inc., Cary, NC, 1989–2021). Subsequent quadratic discrimination function analyses (QDFA) were plotted as canonical correlates to assess the influence of each putative diagnostic miRNA (C-E), depicted as vectors reflecting the contribution of each marker overlaid onto a canonical plot, and to compare their relative association with the single canonical variable. Classification analyses (Group I only) associated with the QDFA were visualized as scatterplot matrices depicting the spatial distribution of positive and negative categories, with 95% confidence interval ellipses estimated and superimposed onto each matrix. These analyses revealed the extent to which individual specimens could be congruently assigned (P<0.05) to their respective IHC test categories (either IHC “positive” or “non-detect”).

### Data Visualization

To determine the distribution of the calculated ΔCt values, we first plotted the number of values for each miRNA in single cycle ΔCt range bins (Fig. 1). Also, as an independent approach to the DFA to assign threshold parameters for miRNA designation, we plotted pair-wise regression analyses of miRNAs and visually partitioned the *a priori* scoring of individual specimens by the NVSL. Specifically, we first conducted logistic regression of ΔCt scatterplots for all six pair-wise combinations among the three potentially diagnostic miRNAs (C, D, and E) to determine correlations and any diagnostic capabilities of miRNAs. Upper and lower thresholds were then designated for the miRNAs in each pair-wise comparison. Then, to illustrate the thresholds, we superimposed a “box” over the pair-wise ΔCt scatterplots and adjusted to capture 100%, 75%, and 50% of the samples identified by IHC as being CWD positive. This partitioning of the data allowed for a projection of minimum and maximum threshold boundaries for assigning miRNAs in the absence of *a priori* IHC data.

**Fig. 1.**
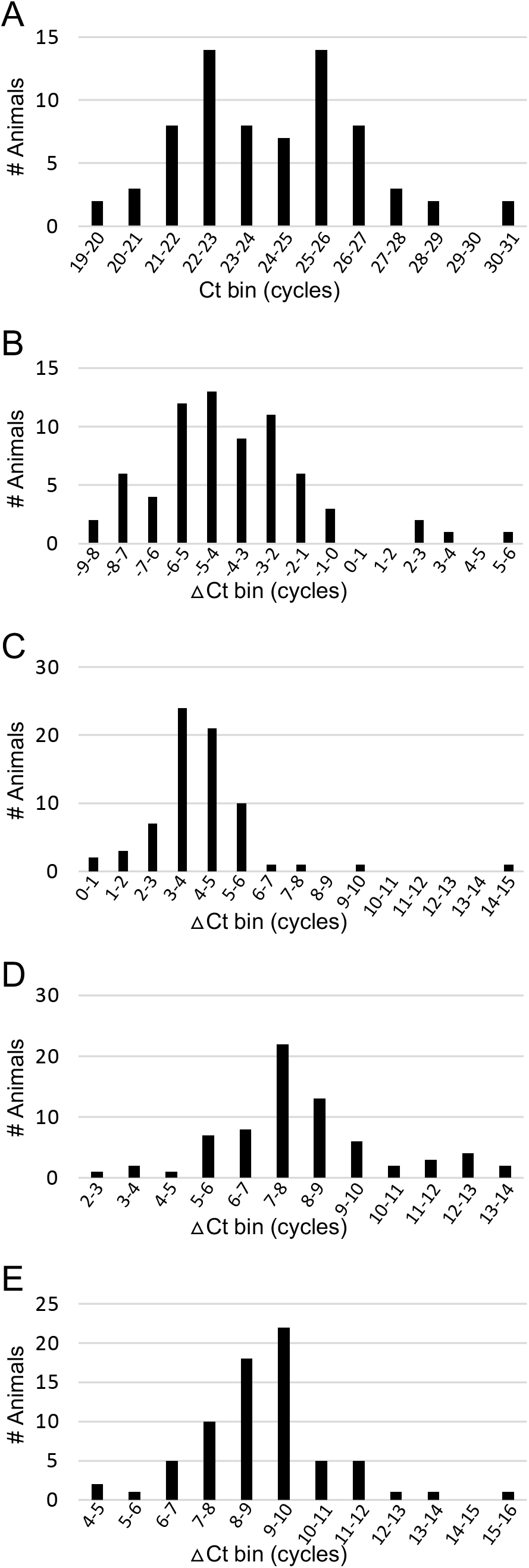
Distribution of Ct and ΔCt values from amplification of reference and putative diagnostic miRNAs. Y-axis indicates number of animals (out of 71 total) in each Ct or ΔCt amplification range bin (# cycles). <u>Panel A</u>: Raw Ct for reference miRNA. <u>Panels B-E</u>: ΔCt calculated by subtracting the reference miRNA Ct from the indicator miRNA Ct to normalize indicator signals for each animal. <u>Panel B</u>: hemolysis indicator miRNA. <u>Panels C-E</u>: putative CWD indicator miRNAs, hereafter designated “diagnostic C,” “diagnostic D,” and “diagnostic E,” respectively. Note the high reference miRNA Ct values returned for a small number of animals, indicative of poor amplification (thus specimens excluded from some subsequent analyses). Note also the mostly negative ΔCt values returned for the hemolysis indicator miRNA, reflecting robust amplification of that marker (raw Ct less than that for the reference miRNA).

## RESULTS

Based on results of a previous study (Slota et al. 2019), we initially selected five miRNAs that might serve as serum biomarkers for pathology-associated changes in animals with CWD. Preliminary quantification experiments identified three of the five miRNA targets (designated C, D, and E) as putative diagnostic markers for prion infection. Among the full set of 93 samples, only 71 returned Ct values for the three putative diagnostic and for putative reference and hemolysis miRNAs. For the 71 samples, raw Ct values for the reference miRNA (designated “A”) and ΔCt for a putative hemolysis miRNA (“B”) and the three diagnostic miRNAs (“C—E”) distributed over relatively large ranges of cycles (Fig. 1), with a small number of specimens requiring an excessive number of cycles for amplification. The distribution of miRNA A (Fig. 1, panel A) appeared generally normal, suggesting its abundance is independent of prion infection. We therefore selected this miRNA to serve as an endogenous reference to normalize diagnostic miRNAs’ amplification values. The small number of specimens that returned Ct’s for the reference miRNA >28 cycles indicated poor (and therefore unreliable, albeit unexplained) amplification, so this result became a possible exclusion criterion for future analyses (see below). Furthermore, analysis of qPCR data for the candidate biomarkers on the 71 samples raised concerns that poor sample quality, evident as moderate to severe hemolysis, might be masking the diagnostic potential of the markers. In addition, the miRNA previously used as a hemolysis marker (Slota et al.; here designated “B”) did not correlate well with hemoglobin concentrations measured either visually (Visual Hemolysis Score, VHS) using the CDC color palette or spectrophotometrically at 414 nm (R^2^ = 0.49), whereas the latter two methods correlated well (NanoDrop vs VHS regression line slope 0.87, R^2^ = 0.92) over the 0-1.2 mg/ml hemoglobin concentration range. We therefore set an upper limit of 1.2 mg/ml hemoglobin concentration as an additional exclusion criterion for degraded samples, which is relaxed in comparison to CDC requirements for human blood tests (less than 1 mg/ml). On applying both exclusion criteria, 38 of the 71 samples examined were too degraded for reliable miRNA quantification, and thus were provisionally excluded from some downstream analyses (32 for excessive hemolysis, six for miRNA A Ct >28).

We next applied discriminant function analysis to the qPCR results for the three putative diagnostic miRNAs to determine the degree of separation in non-linear space among groups of positive and negative individuals (Fig. 2) and to assess congruence between the miRNA biomarker test and diagnoses by IHC (Fig. 3). For all three datasets (Groups I-III), canonical QDFA plots revealed spatial associations among the three diagnostic miRNAs C, D, and E. For Group III (n = 71, Fig. 2. Panel A), miRNA C showed moderate negative association with the Canonical variate 1 (40.1%), whereas miRNA D showed strong positive association (42.3%), and miRNA E showed weak positive association (2.4%). In addition, the length of the vector for miRNA C indicated weaker contribution to variation, whereas length of the vector for miRNA D and E indicated strong contribution to variation. For Group II (n = 39, Fig. 2. Panel B), miRNAs C (9.3%) and E (15.6%) showed weak positive association with the Canonical variate 1, whereas miRNA D showed moderate positive association (25.0%). In addition, the length of the vector for miRNA C indicated weaker contribution to variation, whereas miRNA D and E indicated moderate contribution to variation. For Group I (all exclusion criteria applied; n = 33, Fig. 2. Panel C), miRNA C showed strong positive association with Canonical variate 1 (47.8%), whereas miRNA D showed weak negative association (16.7%) and miRNA E showed weak positive association (24.6%). In addition, the length of the vector for miRNA C and D indicated moderate contribution to variation, whereas length of the vector for miRNA E indicated strong contribution to variation.

**Fig. 2.**
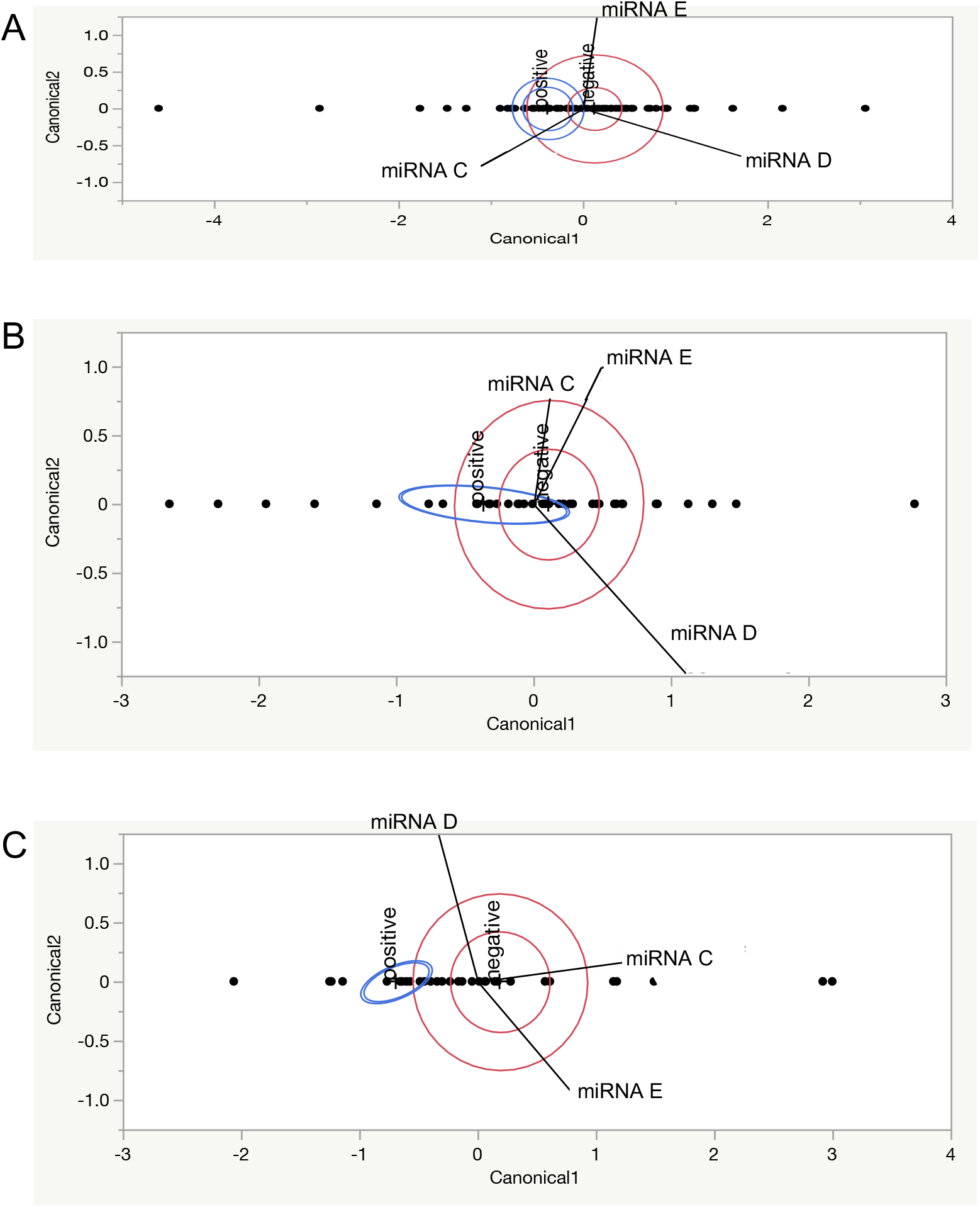
Canonical Plot for QDFA. <u>Panel A</u>: analysis with no data excluded (n=71). <u>Panel B</u>: analysis with data from excessively hemolyzed specimens excluded (n=39). <u>Panel C</u>: analysis with data from hemolyzed and poor reference miRNA amplification excluded (n=33). Note the progressive improvement in resolution of CWD positive specimens with successive application of exclusion criteria.

**Fig. 3.**
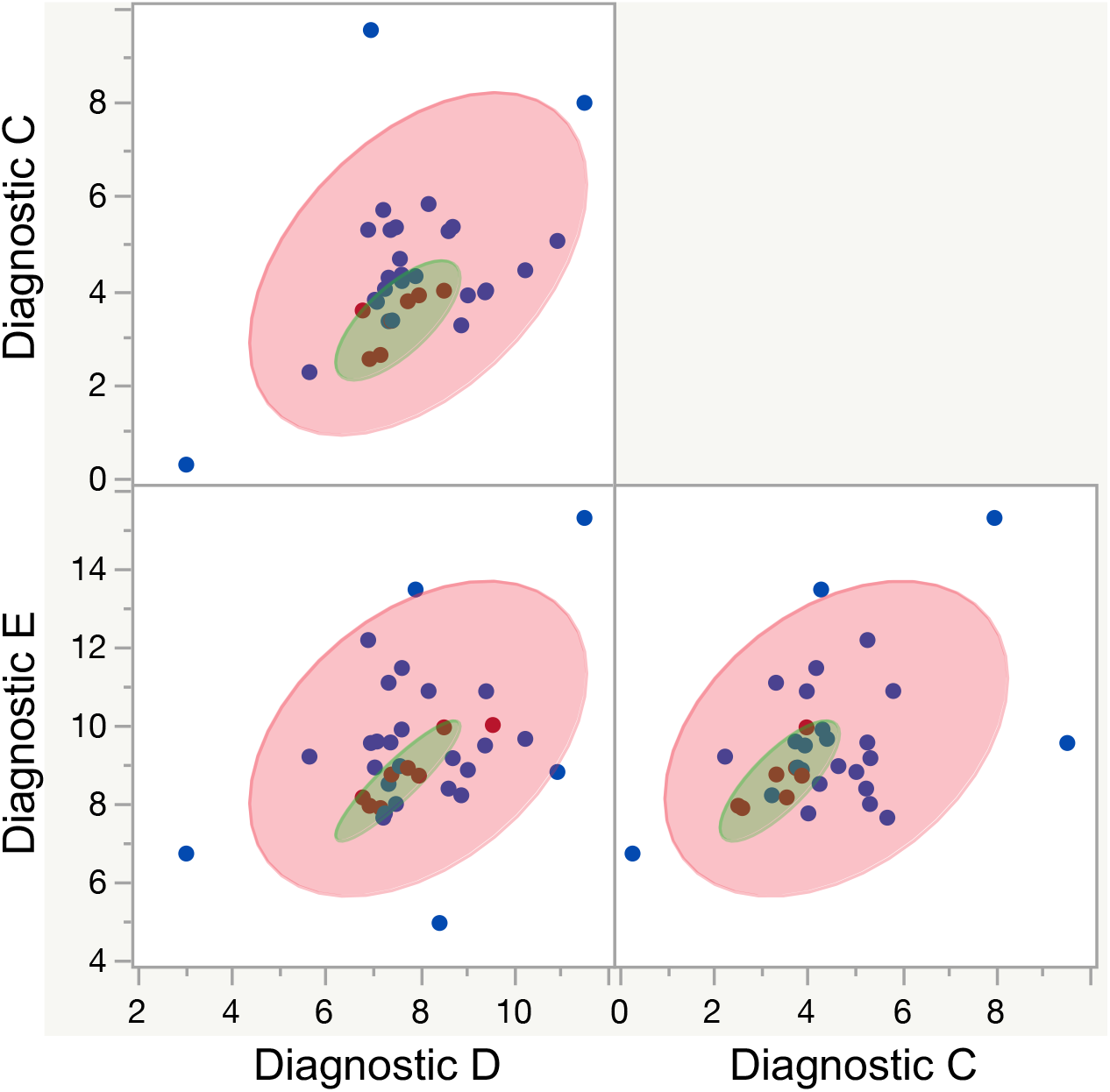
DFA scatterplot matrix. Pairwise comparisons of qPCR data for putative diagnostic miRNAs reveal combined diagnostic potential. Green ellipse depicts 95% confidence interval for specimens that tested CWD positive by IHC (red dots), and the pink ellipse the corresponding interval for negatives (blue dots).

QDFA scatterplot matrices of the ΔCt values for each pair-wise combination of the putative diagnostic miRNAs (C—E, Fig. 3) illustrated the predictive values of all three markers that collectively might constitute a valid miRNA assay for overall classification of specimens from CWD positive and non-detect animals. For Group I (strictest exclusion criteria applied; n = 33), the miRNA assay classified CWD positive and negative animals with 93.9% accuracy overall (Table 1). All seven individuals in Group I identified as positive by IHC also classified positive by miRNA assay (100% sensitivity), and 24 of 26 samples identified as “non-detect” by IHC also classified as negative by miRNA assay (92.3% specificity). For Group II (no exclusion of specimens with poor reference miRNA amplification; n = 39), the miRNA assay classified CWD positive and negative animals with 84.6% accuracy overall. All nine individuals in Group II identified as positive by IHC also classified positive by miRNA assay (100% sensitivity), and 24 of 30 negative samples identified as “non-detect” by IHC also classified negative by miRNA (80% specificity). Finally, for Group III (no exclusion other than specimens previously thawed; n = 71), the overall accuracy of the miRNA assay dropped to 59.2%, with only 14 of 17 samples identified as positive by IHC also classified positive by miRNA assay (82.4% sensitivity), and only 28 of 54 samples identified as negative by IHC also classified negative by mRNA assay (51.9% specificity).

**Table 1.** Classification matrices developed from a discriminant function analysis of the three miRNA markers (C-E) and the three data subsets (Groups I-III). Data exclusion criteria for the Groups are described in the text. Numbers reflect where individual samples were assigned based on ΔCt values. Assignment to “positive” or “negative” categories (a *priori* assumption) was based on an immunohistochemistry (IHC) assay done by the Texas Veterinary, Medical, and Diagnostic Laboratory (TVMDL). Note that among the seven IHC positive animals in Group I, two were positive only in the retropharangeal lymph node (i.e. were negative in obex), indicating that buildup of pathological prion amplicons had not yet reached the nervous system.

| Dataset | Classification by IHC (*a priori*) | DFA Reclassification by miRNA assay |  |
| --- | --- | --- | --- |
|  |  | Negative | Positive |
| Group I, n=33 | Negative (n=26) | 24 | 2 |
|  | Positive (n=7) | 0 | 7 |
| Group II, n=39 | Negative (n=30) | 24 | 6 |
|  | Positive (n=9) | 0 | 9 |
| Group III, n=71 | Negative (n=54) | 28 | 26 |
|  | Positive (n=17) | 3 | 14 |

The miRNA assay’s high overall accuracy as assessed by QDFA reflected its high sensitivity (100% for the Groups I and II datasets), diminished only by lower specificity (“false positives,” i.e. animals that tested positive by miRNA but “non-detect” by IHC). For an independent assessment of the putative diagnostic miRNAs’ utility for detection of CWD infection, individually and in all combinations, and to parse the effects of false positives on the assay’s accuracy, we examined the effects of progressively shrinking the window for identifying positive specimens in pairwise ΔCt scatterplots. Linear regression analyses of each pairwise combination (Fig. 4) revealed positive correlations between each of the three diagnostic miRNAs, with clustering of IHC positive samples at the lower ends of the ΔCt ranges for each miRNA. Despite the positive correlations overall between the putative diagnostic miRNAs’ abundance, however, each of the three appeared to be independently informative, especially for identifying IHC negative specimens. Consequently, to test whether altering thresholds for identifying negative specimens would increase specificity (necessarily at the expense of sensitivity), we adjusted ΔCt range windows/quadrants to capture 100%, 75%, and 50% of the IHC positive samples (Figs. 4A-C), and recalculated specificity and accuracy (“False positives” and “Accuracy” columns, Table 2). The analysis revealed that accuracy increased only marginally upon decreasing ΔCt partitions to capture progressively fewer IHC positive specimens. Given the importance of avoiding false negatives and the relatively low downside risk of a false positive CWD test, as well as the possibility that the test could have identified an infected animal that is presymptomatic, partitioning the data at the higher ΔCt values is most appropriate.

**Fig. 4A.**
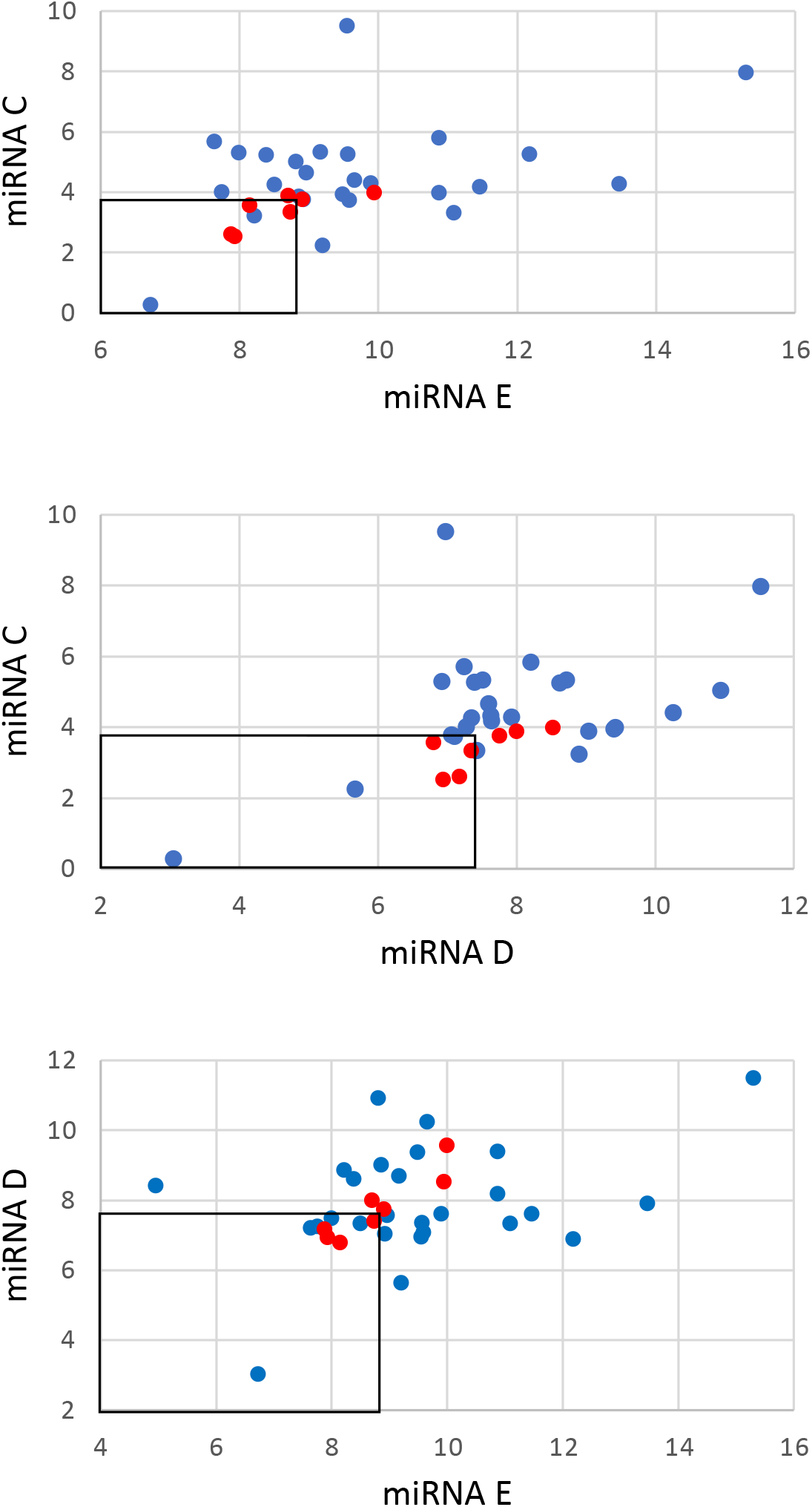
Pairwise regression analyses of three diagnostic miRNA values for Group I with Ct partitions drawn to capture 100% (7/7) of positive samples in the lower left quadrant (boxed). Blue indicates IHC negative and red indicates IHC positive samples. See Table 2 for quantitative summary of these analyses.

**Fig. 4B.**
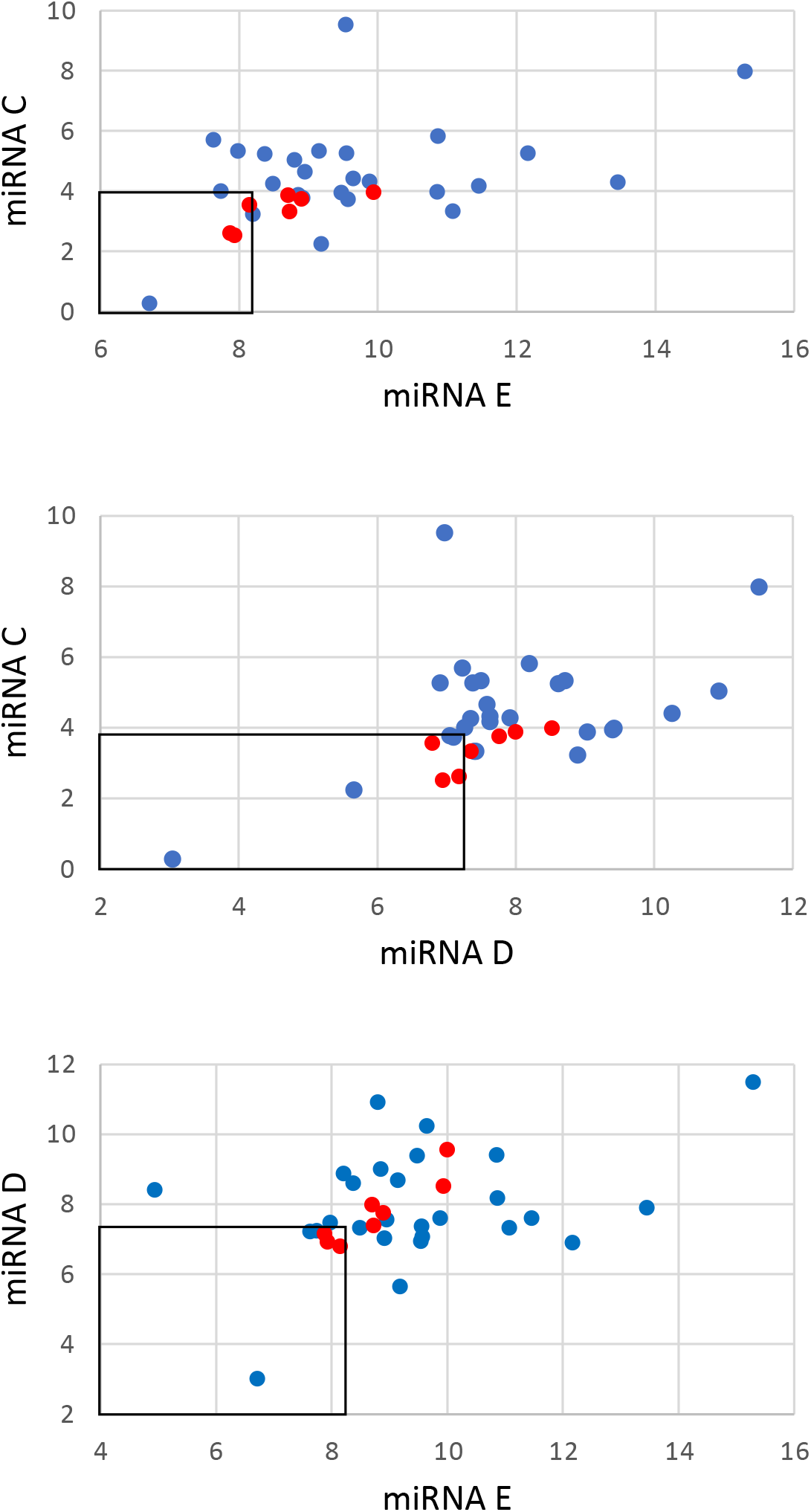
Pairwise regression analyses of three diagnostic miRNApositive samples in the lower left quadrant (boxed). Blue indicates IHC negative and red indicates IHC positive samples. See Table 2 for quantitative summary of these analyses.

**Fig. 4C.**
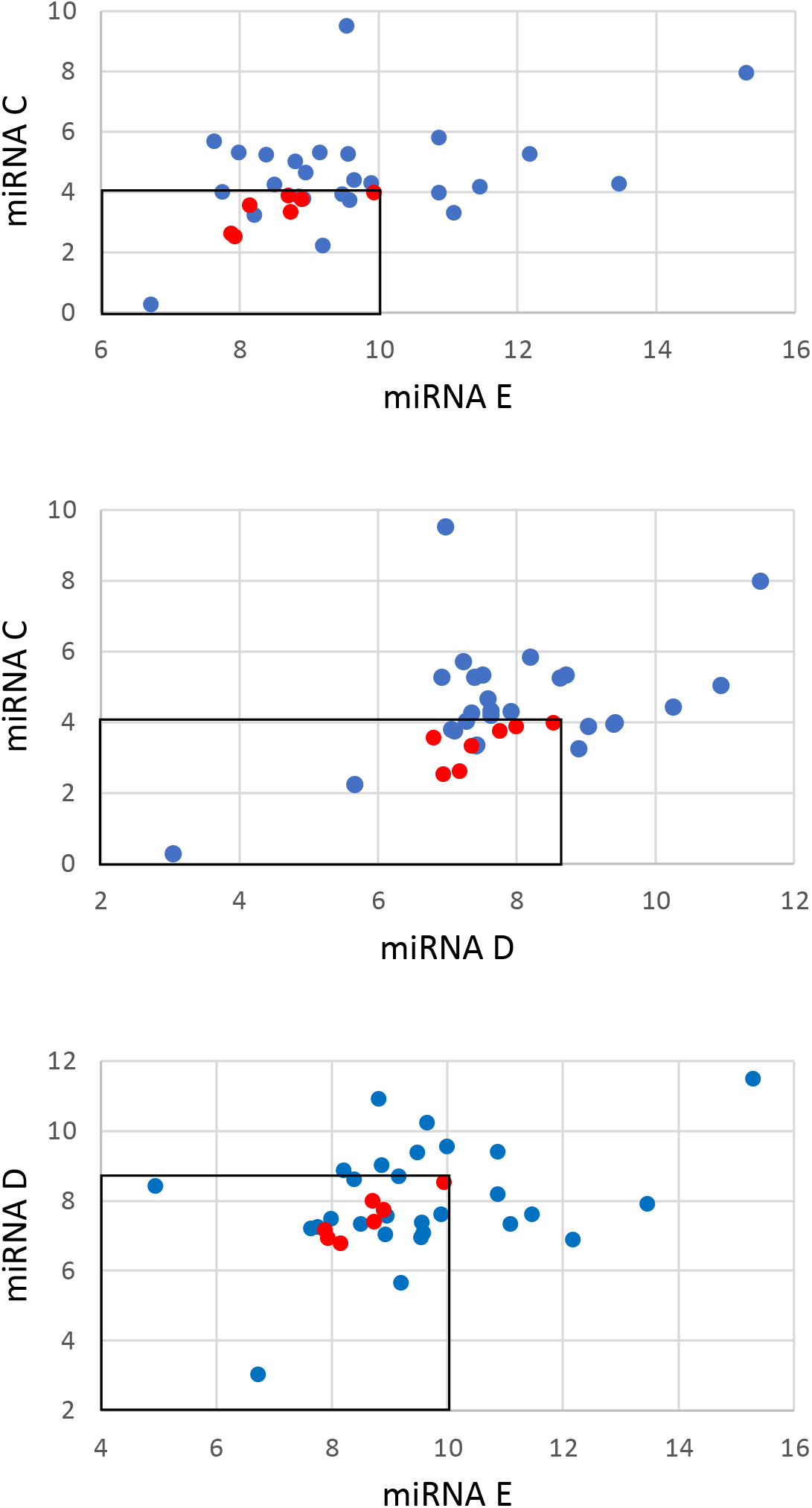
Pairwise regression analyses of three diagnostic miRNApositive samples in the lower left quadrant (boxed). Blue indicates IHC negative and red indicates IHC positive samples. See Table 2 for quantitative summary of these analyses.

**Table 2.**
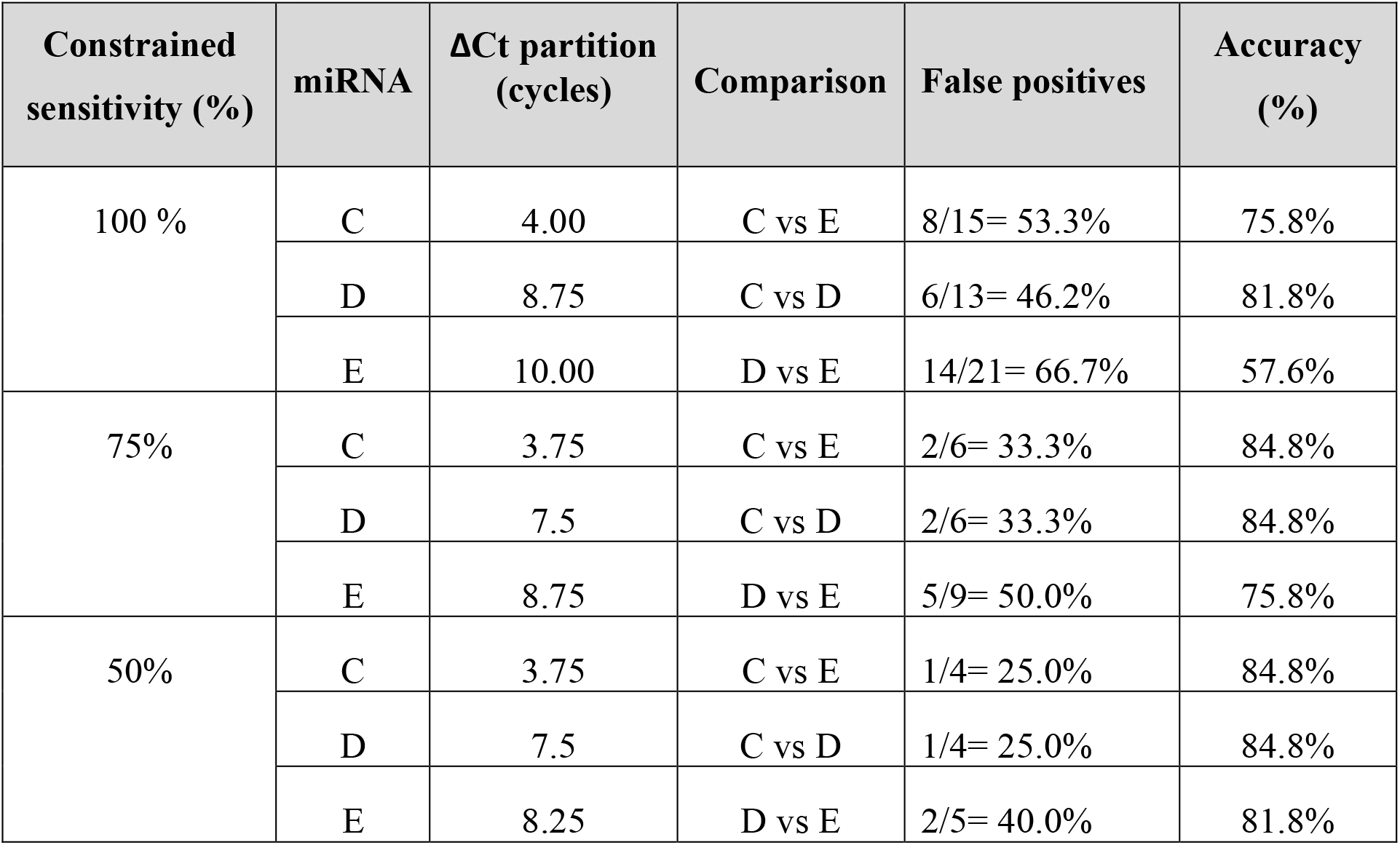
Effect of upper and lower threshold adjustment on miRNA assay accuracy. Constraining the minimal ΔCt diagnostic window in pairwise comparisons of the three miRNA markers (C--E) to 100, 75, or 50% of IHC positive samples (=sensitivity), reflecting 0, 25, and 50% false negatives samples respectively, progressively decreased the percentage of false positives as expected, but at the expense of overall accuracy (two rightmost columns). Number of mismatches was determined by subtracting the number of positive samples from the total number of samples occurring on either side of the threshold boundaries. Note that the ΔCt adjustments variably affected accuracy, with modestly increased accuracy coming at the dramatic expense of sensitivity. Note also that adjusting the capture window required alteration of different miRNAs’ ΔCt partitions, indicating that the three markers were not redundantly informative.

The analyses presented in Figs. 4A-C and summarized in Table 2 also revealed that the three putative diagnostic miRNAs were independently informative. Consequently, to maximize the miRNA test’s accuracy, we combined the three partitions into an aggregate method for identifying positive and negative specimens. Specifically, we identified negatives as those returning ΔCt values greater than any one of the three partitions set to capture positives with 100% sensitivity (Table 2). Collectively, in our admittedly limited dataset, applying these cutoffs for all three miRNAs raised specificity to 92.3%, and accuracy to 93.9%, which is comparable to the accuracy of the other CWD tests currently in use (Table 3).

**Table 3.**
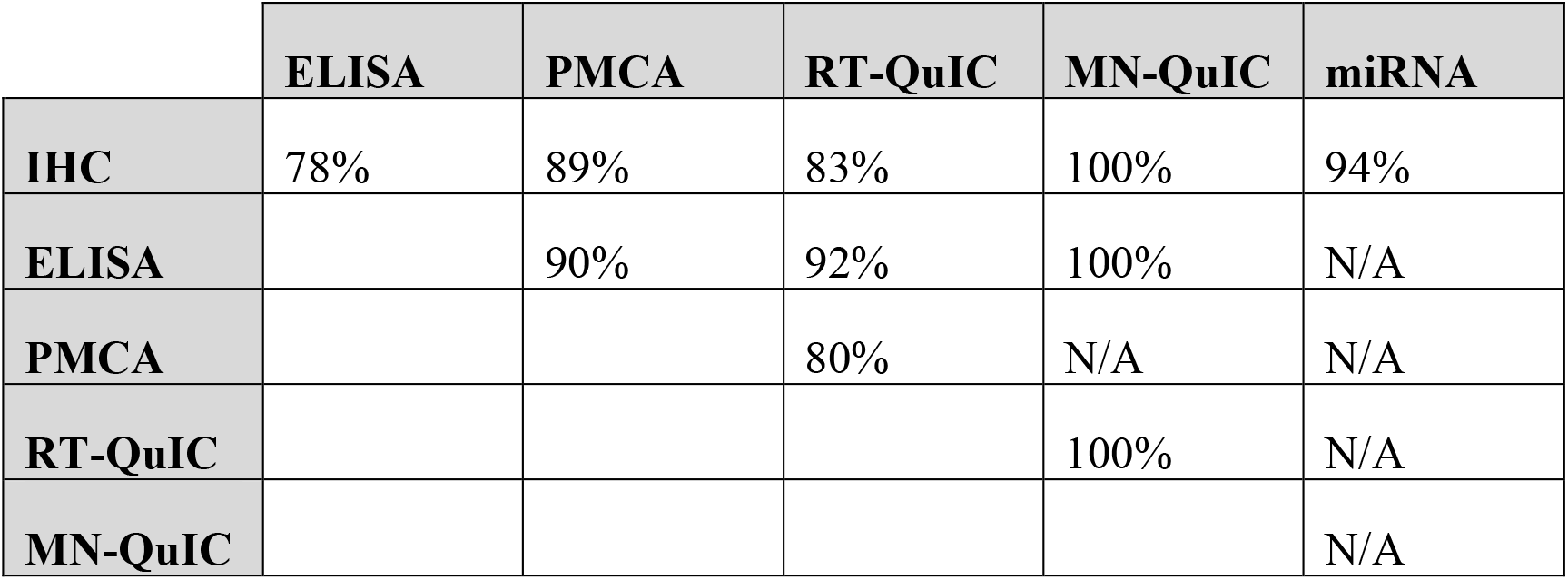
Congruence of current methods for detection of CWD infection. Shown are inter-assay accuracy calculations for Abbreviations are: IHC= <u>I</u>mmuno<u>h</u>isto<u>c</u>hemistry, ELISA= <u>E</u>nzyme-<u>L</u>inked <u>I</u>mmuno<u>s</u>orbent <u>A</u>ssay, PMCA= <u>P</u>rotein <u>M</u>isfolding <u>C</u>yclic <u>A</u>mplification, RT-QuiC= <u>R</u>eal-time <u>Q</u>uaking-Induced <u>C</u>onversion, MN-QuiC= <u>M</u>i<u>n</u>nesota <u>Q</u>uaking-Induced <u>C</u>onversion, miRNA= this study.

|  | **ELISA** | **PMCA** | **RT-QuIC** | **MN-QuIC** | **miRNA** |
| --- | --- | --- | --- | --- | --- |
| **IHC** | 78% | 89% | 83% | 100% | 94% |
| **ELISA** |  | 90% | 92% | 100% | N/A |
| **PMCA** |  |  | 80% | N/A | N/A |
| **RT-QuIC** |  |  |  | 100% | N/A |
| **MN-QuIC** |  |  |  |  | N/A |

## DISCUSSION

Here we described a blood test for prion infection in cervids based upon pathology-associated changes in the abundance of select miRNA biomarkers in serum exosomes. This emerging technology does not depend on detection of prions, unlike all other CWD tests that typically require postmortem tissues and thus cannot be used for routine, antemortem CWD surveillance in herds. Instead, this biomarker test measures physiological sequelae of prion infection, specifically the alteration of cellular miRNA expression and the shedding of exosomes containing those miRNAs into the bloodstream. Although our experiments used postmortem-collected blood (our only option for acquiring research materials from CWD-infected animals), the test itself would best be done using blood obtained non-invasively, such as in the controlled setting of a well-managed, captive herd. Indeed, our sample size was limited due to the necessary exclusion of degraded specimens; the observed sample degradation reflected the difficult collection conditions, specifically the need to prioritize postmortem collection of obex and retropharyngeal lymph node in a mass depopulation setting, not only for experimental purposes but also to comply with federal and state regulations mandating procedures for disposal of CWD-infected herds. Nevertheless, despite obstacles in sample acquisition, we were able to generate a preliminary dataset demonstrating the potential of a highly sensitive, antemortem miRNA CWD test.

Initial hemolysis estimates and reference miRNA amplification sorted sera from 71 animals into three subsets (Groups I, II, and III) based on serum degradation (minimal, moderate, and high; respectively). Comparative histogram analyses (Fig. 1) indicated that reference miRNA A was unaffected by prion association, confirming its utility as an endogenous control for normalization. Pairwise regression analyses between all proposed diagnostic miRNAs revealed their individual utility for detecting prion infections, and setting a threshold ΔCt value for each miRNA increased specificity. A subsequent QDFA with corresponding scatterplots (Fig. 2-3) of Groups I, II, and III demonstrated progressive loss of test accuracy with increasing specimen degradation. For Group I, the DFA classified positive and negative samples with 92.3% accuracy compared to the IHC designation performed by NVSL, but as sample quality declined (Groups II and III), the QDFA classified samples with lower accuracy (84.6% and 59.15%, respectively). Furthermore, in Group III the miRNA assay began to detect false negatives (positive by IHC but called negative by miRNA), which would be considered unacceptable for a CWD detection test. Nevertheless, the QDFA did show that changes in diagnostic miRNA abundance were able to discriminate between positive and negative samples classified by IHC. For the non-degraded sample set (Group I) the only discrepancies between testing methods were false positive results (two animals called positive by miRNA assay that tested non-detect by IHC). It is formally possible that this disagreement between testing methods reflects ability of the miRNA assay to detect a pre-clinical infection. Indeed, the miRNA assay did identify two animals (out of seven in Group I) that tested positive by IHC only in retropharangeal lymph nodes, indicative of possible early stage infections. Nevertheless, further study will be required to clarify the relationship between miRNA content in exosomes and progression of CWD pathophysiology.

As a companion system to the QDFA, threshold value maxima for each miRNA, assigned by visual partitioning, also reliably identified of positive and negative samples (as determined by IHC). Among the three threshold parameters examined (100%, 75%, and 50%), the 100% and 75% categories established spatial boundaries with maximal inclusion of positive samples, while minimizing test discrepancies, both false positives and false negatives. The modestly improved accuracy achieved by downward adjustment of the ΔCt partitions for capturing positives in each pairwise comparison necessarily came at dramatic expense of sensitivity, which for a CWD test is an unacceptable tradeoff. We therefore propose that setting individual partitions to capture all positive specimens, applying a hierarchical diagnostic protocol with miRNA C as the most strongly discriminating component, and collectively applying data from all three diagnostic markers would produce an optimal, miRNA-based, high throughput assay for CWD infection.

Sample quality influenced amplification of miRNAs, demonstrating that the timing of postmortem sampling may be critical for the reliability of this assay. Observed, excessive hemolysis likely reflects sample degradation that may have perturbed quality of miRNA templates, resulting in unreliable amplification of miRNAs and subsequent poor congruence between miRNA abundance and CWD in our Group III dataset. Among the three subsets of data, Group I yielded only two discrepancies, where the IHC result was negative and miRNA result was positive. Group II yielded six discrepancies where the IHC result was negative and miRNA result was positive. But Group III yielded both types of discrepancy, with 26 positives by miRNA testing negative by IHC and three negatives by miRNA testing positive by IHC. For samples not eliminated by the exclusion criteria, few disparities will occured, and then only in the direction of a false-positive result. As samples became more degraded (Group III), the number of disparities (n = 29) between IHC-negative: miRNA-positive (n = 26) and IHC-positive: miRNA-negative (n = 3) increased, confirming that degradation decreased accuracy. Thus, applying hemolysis and miRNA integrity exclusion criteria proved important for maximizing the accuracy of the assay, stressing the likely value of antemortem sample collection. Indeed, acquiring blood during routine animal husbandry (captive deer) or immediately after death (wild deer) appears to be essential for this test to yield reliable results.

For the minimal serum degradation group, miRNA C appeared to have the strongest association with the canonical variant 1 (Fig. 2) and to have contributed the greatest weight in classifying positive and negative groups in the QDFA analysis. Comparatively, miRNA D and E were less informative. Although it appears that miRNA C was strongly diagnostic, it is premature to speculate whether it alone might serve as a reliable marker for CWD, or if the additive combination of miRNAs were synergistic in their classification among positive and negative individuals. miRNA C is less influential on subsets II and III (n = 39 and n = 71). With those two subsets, miRNA D appears to have a moderate degree of association with canonical variant 1. In estimating the minimal and maximal threshold boundaries for each of the three miRNAs, miRNA C had a narrower threshold range than miRNA D and E. This observation indicates that miRNA C possesses a narrow range of variation when classifying samples as either positive or negative. For the remaining two miRNAs, miRNA D is slightly more discriminant than miRNA E when classifying samples as either positive or negative. The above observations are consistent regardless of sample quality in the dataset.

The results by this miRNA assay compared favorably to the IHC test provided by TVMDL with a range of 86-93% accuracy when using samples with minimal to moderate serum degradation. These results (Table 3) are similar to, or in some cases, exceed comparisons of PMCA, Rt-QuiC, and Immuno-QuiC to IHC and ELISA scores as reported in previous studies (80-82%; Saborio et al. 2001; Soto et al. 2002; Castilla et al. 2005; Hibler et al. 2003; Keane et al. 2008, 2009; Daus et al., 2011; Saa et al 2006; Thomsen et al. 2012).

Given the complementary results, ease, and speed at which miRNAs assays can be completed, it appears miRNA techniques may offer advantages for high throughput CWD testing. Generally, a sample can pass through the entire process (sample process and RNA isolation to data processing) in under 12 hours. In addition, Groups I and II, the disparities between the miRNA assay and IHC results could be interpreted as early detection of biomarkers indicative of CWD pathology in presymptomatic or IHC non-detect animals, which could be valuable for surveillance. However, given that the miRNA assay does not directly detect prions or plaques, it is not intended to serve as a replacement for IHC test but instead as a complementary tool.

Although this study was conducted on post-mortem blood samples, the long-term goal is to develop an antemortem blood assay that, subject to federal and varying state regulations, could prove useful as an early-detection/rapid response strategy for CWD surveillance, to both the scientific community and the private sector. Application of this miRNA test to antemortem sampling could enable: 1) a high throughput capability and rapid return of results (same day for a single sample assay, in most cases); 2) repeated testing of the same individual with minimal handling and stress; 3) where permitted by current or future regulations, selective culling of known positive animals rather than destruction of all individuals connected to diseased animals or a contaminated geographic location; 4) continual monitoring or periodic testing in geographic locations or facilities at risk of having positive individuals; 5) cooperation among landowners, deer breeders, and governmental agencies for management and surveillance for CWD in deer herds, and for possible establishment of regulations permitting sale and relocation of animals that test negative; 6) monitoring of wild populations by state and federal agencies; 7) early detection (relative to IHC) of presymptomatic animals; and 8) research aimed at modeling the spread of disease into populations with varying genetic susceptibilities, and at understanding the association between progression of overt disease (manifestation of wasting and cognitive decline symptoms) and its underlying pathophysiology (e.g. perturbation of miRNA profiles).

Difficulty in getting non-degraded specimens hampered our efforts on this study. Indeed, considering the generally poor quality of available specimens (an inevitable consequence of post-mortem acquisition under the very difficult field conditions of a depopulation event) and resultant need to limit our analysis to only those samples that met defined exclusion criteria, our findings provide some optimism for future development and implementation of miRNA-based test for CWD infection. Unfortunately, our data provide little insight into how sample collection 1-12 hours post-euthanasia affected degradation of miRNAs, though other studies identify hemolysis as a strong indicator of specimen degradation. Regardless, acquiring samples under better field conditions so more of them meet our exclusion criteria will greatly aid in the ongoing development of miRNA-based tests for high throughput screening of wild and captive animals. Such tests will provide the tools necessary to track disease progression in captive herds and thereby define miRNA changes indicative of different stages of infection, and in turn define and potentially avoid circumstances that might lead to depopulation of entire herds. Finally, this method may be extended to different sample types (e.g. skeletal muscle) that applies postmortem sampling to provide more convenient testing for the hunting industry.

## ACKNOWLEDGMENTS

We thank M. Lockwood and H. Reed of the Texas of Parks and Wildlife Department for sample acquisition. We thank W. Conway and C. Ramsey for helping with sample logistics and comments on an earlier draft of the manuscript.

## AUTHOR CONTRIBUTIONS

Robert D. Bradley - PI, project organization, communication with many, laboratory supervision (work done in his laboratory), troubleshooting, drafting and editing the manuscript

Emma K. Roberts- organization, laboratory work, troubleshooting, drafting and editing the manuscript

Asha Worsham - laboratory work, data analysis and interpretation, drafting and editing the manuscript

Megan Ashton - laboratory work, data analysis and interpretation, drafting and editing the manuscript

Emily A. Wright - laboratory work, troubleshooting, editing the manuscript

Ned Saleh – data analysis and resources (experimental protocols, data analysis tools)

Daniel M. Hardy - PI, organization, supervision (work done in his laboratory), troubleshooting, data analysis and interpretation, drafting and editing the manuscript

